# Evaluating causal associations between previously reported risk factors and epithelial ovarian cancer: a Mendelian randomization analysis

**DOI:** 10.1101/472696

**Authors:** James Yarmolinsky, Caroline L Relton, Artitaya Lophatananon, Kenneth Muir, Usha Menon, Aleksandra Gentry-Maharaj, Axel Walther, Jie Zheng, Peter Fasching, Wei Zheng, Woo Yin Ling, Jenny Chang-Claude, Sue K Park, Byoung-Gie Kim, Ji-Yeob Choi, Boyoung Park, George Davey Smith, Richard M Martin, Sarah J Lewis

## Abstract

**Background:** Various modifiable risk factors have been associated with epithelial ovarian cancer risk in observational epidemiological studies. However, the causal nature of the risk factors reported, and thus their suitability as effective intervention targets, is unclear given the susceptibility of conventional observational designs to residual confounding and reverse causation. Mendelian randomization uses genetic variants as proxies for modifiable risk factors to strengthen causal inference in observational studies. We used Mendelian randomization to evaluate the causal role of 13 previously reported risk factors (reproductive, anthropometric, clinical, lifestyle, and molecular factors) in overall and histotype-specific epithelial ovarian cancer in up to 25,509 case subjects and 40,941 controls in the Ovarian Cancer Association Consortium.

**Methods and Findings:** Genetic instruments to proxy 13 risk factors were constructed by identifying single nucleotide polymorphisms (SNPs) robustly (*P*<5×10^−8^) and independently associated with each respective risk factor in previously reported genome-wide association studies. SNPs were combined into multi-allelic inverse-variance weighted fixed or random-effects models to generate causal estimates. Three complementary sensitivity analyses were performed to examine violations of Mendelian randomization assumptions: MR-Egger regression and weighted median and mode estimators. A Bonferroni-corrected *P*-value threshold was used to establish “strong evidence” (*P*<0.0038) and “suggestive evidence” (0.0038<*P*<0.05) for associations.

In Mendelian randomization analyses, there was strong or suggestive evidence that 9 of 13 risk factors had a causal effect on overall or histotype-specific epithelial ovarian cancer. There was strong evidence that genetic liability to endometriosis increased risk of epithelial ovarian cancer (OR per log odds higher liability:1.27, 95% CI: 1.16-1.40; *P*=6.94×10^−7^) and suggestive evidence that lifetime smoking exposure increased risk of epithelial ovarian cancer (OR per unit increase in smoking score:1.36, 95% CI: 1.04-1.78; *P*=0.02). In histotype-stratified analyses, the strongest associations found were between: height and clear cell carcinoma (OR per SD increase:1.36, 95% CI: 1.15-1.61; *P*=0.0003); age at natural menopause and endometrioid carcinoma (OR per year later onset:1.09, 95% CI: 1.02-1.16; *P*=0.007); and genetic liability to polycystic ovary syndrome and endometrioid carcinoma (OR per log odds higher liability:0.74, 95% CI:0.62-0.90; *P*=0.002). There was little evidence for an effect of genetic liability to type 2 diabetes, parity, or circulating levels of 25-hydroxyvitamin D and sex hormone-binding globulin on ovarian cancer or its subtypes. The primary limitations of this analysis include: modest statistical power for analyses of risk factors in relation to some less common ovarian cancer histotypes (low grade serous, mucinous, and clear cell carcinomas), the inability to directly examine the causal effects of some ovarian cancer risk factors that did not have robust genetic variants available to serve as proxies (e.g., oral contraceptives, hormone replacement therapy), and the assumption of linear relationships between risk factors and ovarian cancer risk.

**Conclusions:** Our comprehensive examination of possible etiological drivers of ovarian carcinogenesis using germline genetic variants to proxy risk factors supports a causal role for few of these factors in epithelial ovarian cancer and suggests distinct etiologies across histotypes. The identification of novel modifiable risk factors remains an important priority for the control of epithelial ovarian cancer.

## Introduction

Ovarian cancer is the second most common gynecological cancer in the USA and Western Europe and accounts for more deaths than all other gynecological cancers combined ^1,2^. The prognosis for ovarian cancer is generally poor because women typically present with advanced disease due to the non-specific nature of symptoms and because of the lack of established screening tests ^3-5^. Given the limited success of secondary prevention strategies and the sporadic nature of 90% of cases, primary prevention of ovarian cancer may serve as an important vehicle for disease control ^6^. However, few modifiable risk factors have consistently been linked to ovarian cancer in observational epidemiological studies and most previous studies have failed to stratify analyses across clinically distinct histotypes ^7-10^. Further, the causal nature of the risk factors reported, and thus their suitability as effective intervention targets, is unclear given the susceptibility of conventional observational designs to residual confounding and reverse causation.

Mendelian randomization (MR) is an analytical approach that uses germline genetic variants as instruments (“proxies”) for potentially modifiable risk factors, to examine the causal effects of these factors on disease outcomes in observational settings ^11,12^. Since germline genetic variants are randomly assorted at meiosis, MR analyses should be less prone to confounding by lifestyle and environmental factors than conventional observational studies. Further, since germline genetic variants are fixed at conception and cannot be influenced by subsequent disease processes, MR analyses are not subject to reverse causation bias. An additional advantage of MR is that it can be implemented using summary genetic association data from two independent samples, representing: a) the genetic variant-risk factor associations; and b) the genetic variant-outcome associations (“two-sample Mendelian randomization”). This provides an efficient and statistically robust method of appraising causal relationships between risk factors and disease outcomes.

Given the current poor understanding of the etiology of epithelial ovarian cancer (EOC), a two-sample Mendelian randomization analysis was performed to evaluate the causal effects of 13 previously reported factors with risk of overall and histotype-specific EOC.

## Methods

### Ovarian cancer population

Summary genetic association data were obtained on 25,509 women with EOC and 40,941 controls of European descent. These women had been genotyped using the Illumina Custom Infinium array (OncoArray) as part of the Ovarian Cancer Association Consortium (OCAC) genome-wide association study (GWAS) ^13,14^. The data included the following invasive ovarian cancer histotypes: high grade serous carcinoma (n=13,037), low grade serous carcinoma (n=1,012), mucinous carcinoma (n=1,417), endometrioid carcinoma (n=2,810), and clear cell carcinoma (n=1,366). Analyses were also performed for low malignant potential tumors (n=3,103) which included 1,954 serous and 1,140 mucinous tumors. Invasive histotypes classified as “other” (n=2,764 cases) were included in analyses for overall epithelial ovarian cancer but were not assessed separately. Ethical approval from relevant research ethics committees was obtained for all studies in OCAC and written, informed consent was obtained from all participants in these studies. Further details about the OCAC study and OncoArray analyses are available in **Supplemental Materials**.

### Identification of previously reported risk factors and instrument selection

Previously reported risk factors for EOC were identified from a literature review of narrative and systematic review articles summarizing findings from observational epidemiological studies using PubMed and Web of Science ^15-20^ and through consultation with the Cancer Research UK website and the World Cancer Research Fund/American Institute for Cancer Research Ovarian Cancer 2014 Report (accessed on 02/10/2017). Genetic instruments were then identified for these risk factors by consulting the preprint server bioRxiv (http://www.biorxiv.org/) and two catalogues of summary GWAS data: the NHGRI-EBI (National Human Genome Research Institute - European Bioinformatics Institute) GWAS catalogue and MR-Base ^21,22^. The complete PubMed and Web of Science search strategies and instrument selection criteria are presented in **Supplementary Materials** and **Extended Methods**, respectively.

In total, 13 risk factors with a suitable genetic instrument were included in the analysis: four reproductive factors (age at menarche, age at natural menopause, parity, and genetic liability to twin pregnancy)^23-26^, two anthropometric traits (body mass index, height)^27,28^, three clinical factors (genetic liabilities to type 2 diabetes, endometriosis, and polycystic ovary syndrome) ^29-31^, two lifestyle factors (lifetime smoking exposure, circulating 25-hydroxyvitamin D)^32,33^, and two molecular risk factors (C-reactive protein, sex hormone-binding globulin) ^34,35^. Lifetime smoking exposure is a composite score that captures smoking duration, heaviness, and cessation among both smokers and non-smokers. A step-by-step overview of risk factor inclusion along with a flow-chart of these processes and a list of all risk factors ascertained for inclusion are presented in **Supplementary Materials** and **Supplementary Figure 1**.

### Statistical analyses

The use of genetic instruments for potentially modifiable exposures in an MR framework allows for unbiased causal effects of risk factors on disease outcomes to be estimated if: i) the genetic instrument (typically, one or more independent single-nucleotide polymorphisms [SNPs]) is robustly associated with the risk factor of interest; ii) the instrument is not associated with any confounding factor(s) of the association between the risk factor and outcome; and iii) there is no pathway through which an instrument influences an outcome except through the risk factor (“exclusion restriction criterion”).

Estimates of the proportion of variance in each risk factor explained by the genetic instruments (R^2^) and the strength of the association between the genetic instruments and risk factors (F-statistics) were generated using methods previously described ^36^. F-statistics can be used to examine whether results are likely to be influenced by weak instrument bias: i.e., reduced statistical power to reject the null hypothesis when an instrument explains a limited proportion of the variance in a risk factor.

For risk factors with only one SNP as an instrument, the Wald ratio was used to generate effect estimates, and the delta method was used to approximate standard errors ^37^; for risk factors with two or three SNPs as instruments, inverse-variance weighted (IVW) fixed effects models were used; and for risk factors with greater than three SNPs, IVW multiplicative random effects models (allowing overdispersion in the model) were used ^38^. The combination of multiple SNPs into a multi-allelic IVW model increases the proportion of variance in a risk factor explained by an instrument. Causal estimates from these models represent a weighted average of individual Wald ratios across SNPs using inverse-variance weighted meta-analysis. To account for multiple testing, a Bonferroni correction was used to establish *P*-value thresholds for “strong evidence” (*P*<0.0038) (false positive rate=0.05/13 risk factors) and “suggestive evidence” (0.0038<*P*<0.05) for reported associations.

When using genetic instruments, there is potential for horizontal pleiotropy - when a genetic variant has an effect on two or more traits through independent biological pathways, a violation of the third IV assumption. This was examined by performing three complementary sensitivity analyses, each of which makes different assumptions about the underlying nature of horizontal pleiotropy: i) MR-Egger regression (intercept and slope terms);^39^ii) a weighted median estimator ^40^ when there were, at minimum, three SNPs in an instrument; and iii) a weighted mode estimator ^41^ when there were, at minimum, five SNPs in an instrument. Additionally, leave-one-out permutation analyses were performed to examine whether any results were driven by individual SNPs in IVW models. Lastly, Steiger filtering was employed to orient the direction of causal relationships between presumed risk factors and outcomes for some analyses^42^. This method compares the proportion of risk factor and outcome variance explained by SNPs used as instruments to help establish whether SNPs associated with both risk factors and outcomes primarily represent either: 1) a direct association of a SNP on a risk factor which then influences levels of an outcome or 2) a direct association of a SNP on an outcome which then influences levels of a risk factor. Extended descriptions of these sensitivity analyses, along with their assumptions are provided in the **Extended Methods** section.

All statistical analyses were performed using R version 3.3.1.

## Results

Across the 13 risk factors that we examined, F-statistics for their respective genetic instruments ranged from 4 to 423, with 12 of 13 risk factors having a value of F≥24. These statistics suggest that most analyses were unlikely to suffer from weak instrument bias. For each risk factor, the number of SNPs included in the genetic instrument, along with R^2^ and F-statistics for the instrument, are provided in **Supplementary Table 1**. Complete primary and sensitivity analyses for all risk factors categorized by ovarian cancer histotype are presented in **Supplementary Tables 2-6**.

### Reproductive factors

In IVW models, there was suggestive evidence for an effect of earlier age at menarche on risk of overall EOC (OR per year earlier onset: 1.07,95% CI:1.00-1.14;*P*=0.046) and endometrioid carcinoma (OR:1.19,95% CI:1.05-1.36;*P*=0.008) (**Figure 1**). However, there was evidence that horizontal pleiotropy was likely biasing the IVW estimate for EOC. This is because the effect estimate attenuated toward the null when employing MR-Egger regression (OR:1.00,95% CI:0.89-1.13) and a weighted median estimator (OR:1.01,95% CI:0.92-1.10) and moved in a protective direction when using a weighted mode estimator (OR:0.98,95% CI:0.25-3.84). In contrast to EOC, the effect of age at menarche on endometrioid carcinoma was robust to MR-Egger, weighted median, weighted mode estimates, and leave-one-out analyses (**Supplementary Table 2**).

There was suggestive evidence for an effect of later age at natural menopause on risk of endometrioid carcinoma (OR per year later onset:1.09,95% CI:1.02-1.16;*P*=0.007), which was consistent in sensitivity analyses examining horizontal pleiotropy. While there was little evidence of an effect of age at natural menopause on clear cell carcinoma in IVW models (OR:1.05,95% CI:0.96-1.14;*P*=0.29), the association strengthened when employing MR-Egger (OR:1.26,95% CI:1.05-1.52), weighted median (OR:1.11,95% CI:0.99-1.25), and weighted mode estimators (OR:1.16,95% CI:1.02-1.31), suggesting horizontal pleiotropy in the IVW model. There was also suggestive evidence for an effect of genetic liability to twin births on clear cell carcinoma (OR:1.78,95% CI:1.05-3.03;*P*=0.03) which was robust to sensitivity analyses examining horizontal pleiotropy.

In parity analyses, effect estimates were in a protective direction for five of seven ovarian cancer outcomes but were imprecisely estimated with 95% confidence intervals crossing the null line (**Supplementary Table 2**).

### Anthropometric traits

There was strong evidence for an effect of body mass index (BMI) on overall EOC (OR per 1-standard deviation (SD; 4.6 kg/m^2^) increase:1.23,95% CI:1.07-1.42;*P*=0.003) (**Figure 2**). Though there was little evidence for horizontal pleiotropy when performing MR-Egger (OR:1.32,95% CI:0.88-1.99), inconsistency of effect estimates across weighted median (OR:1.14,95% CI:0.93-1.40) and weighted mode (OR:1.05,95% CI:0.75-1.51) approaches suggested potential violations of the IV assumptions.

In IVW models, there was suggestive evidence for an effect of BMI on high grade serous carcinoma (OR:1.26,95% CI:1.06-1.50;*P*=0.01), endometrioid carcinoma (OR:1.48,95% CI:1.07-2.06;*P*=0.02), and low malignant potential tumors (OR:1.39,95%CI:1.04-1.85;*P*=0.03) but not on other histotypes. However, there was evidence that horizontal pleiotropy was likely biasing the IVW estimate for high grade serous carcinoma: the effect estimate was attenuated when performing MR-Egger (OR:1.05,95% CI:0.63-1.75) and was inconsistent when employing weighted median (OR:1.17,95% CI:0.91-1.50) and weighted mode (OR:0.95,95% CI:0.53-1.35) estimators. Likewise, there was some inconsistency of effect estimates across sensitivity analyses for low malignant potential tumors, with a modest attenuation of the effect estimate observed when employing a weighted mode estimator (OR:1.17,95%CI:0.55-2.49). In contrast, the effect of BMI on endometrioid carcinoma was also seen across sensitivity analyses using MR-Egger, weighted median, and weighted mode estimators, and in leave-one-out analyses (**Supplementary Table 3**).

There was strong evidence for an effect of height on clear cell carcinoma (OR per 1-SD (6.3 cm) increase:1.36,95% CI:1.15-1.61;*P*=0.0003), but not on other histotypes. This finding was robust to various sensitivity analyses.

### Clinical factors

There was strong evidence for an effect of genetic liability to endometriosis on EOC (per unit log odds higher liability to endometriosis: OR 1.27,95% CI:1.16-1.40;*P*=6.94×10^−7^) and clear cell carcinoma (OR:2.69,95% CI:1.88-3.86, *P*=7.39×10^−8^) and suggestive evidence for an effect on endometrioid carcinoma (OR:1.37,95% CI:1.10-1.69;*P*=0.004), low malignant potential tumors (OR:1.33,95%CI:1.09-1.63;*P*=0.006), and high grade serous carcinoma (OR:1.17,95% CI:1.04-1.31;*P*=0.007) (**Figure 3**). Findings for overall and clear cell carcinoma were also seen in sensitivity analyses examining horizontal pleiotropy, whereas inconsistent effect estimates for endometrioid carcinoma, low malignant potential tumors, and high grade serous carcinoma across these sensitivity analyses suggested violations of IV assumptions (**Supplementary Table 4**). Analyses employing Steiger filtering provided strong evidence that the causal direction between genetic liability to endometriosis and EOC was from the former to the latter (*P*<10^−10^), whereas the causal direction could not be clearly established for clear cell carcinoma analyses (*P*<0.10).

There was strong evidence for an inverse effect of genetic liability to polycystic ovary syndrome (PCOS) on endometrioid carcinoma (OR per unit log odds higher liability to PCOS:0.74,95% CI:0.62-0.90;*P*=0.002), which was robust to sensitivity analyses. In contrast, suggestive evidence for an effect of PCOS with low grade serous carcinoma (OR:1.33,95% CI:1.01-1.74;*P*=0.04) in IVW models was not seen across all sensitivity analyses examining horizontal pleiotropy. There was little evidence of an effect of genetic liability to type 2 diabetes on overall or histotype-specific ovarian cancer.

### Lifestyle factors

There was suggestive evidence for an effect of lifetime smoking exposure on EOC (OR per unit increase in smoking score:1.36,95% CI:1.04-1.78,*P*=0.02) (**Figure 4**). In histotype-specific analyses, there was also a suggestive association for an effect of smoking on high grade serous carcinoma (OR:1.44,95% CI:1.05-1.98;*P*=0.02) but little association with other subtypes. The smoking findings for epithelial ovarian cancer and high grade serous carcinoma were robust to horizontal pleiotropy sensitivity analyses (**Supplementary Table 5**). There was no strong or suggestive evidence that circulating 25-hydroxyvitamin D influenced overall or histotype-specific ovarian cancer.

### Molecular risk factors

There was suggestive evidence for an inverse effect of C-reactive protein (CRP) on endometrioid carcinoma (OR per unit increase in natural log CRP:0.90,95% CI:0.82-1.00;*P*=0.049) (**Figure 5**). This association was robust to sensitivity analyses using MR-Egger, weighted median, and weighted mode methods in addition to using a restricted CRP instrument (exclusively using 4 SNPs in *CRP*): OR:0.72,95% CI:0.42-1.22;*P*=0.14 (**Supplementary Table 6**). CRP was not clearly associated with other histotypes assessed. There was no strong or suggestive evidence for an effect of sex hormone-binding globulin on ovarian cancer risk.

## Discussion

This Mendelian randomization analysis of up to 66,450 women supports causal effects of liability to endometriosis and lifetime smoking exposure in epithelial ovarian cancer risk but found little evidence for causal roles of eleven previously reported risk factors in ovarian carcinogenesis. In histotype-stratified analyses, there was strong or suggestive evidence of effects of ages at menarche and natural menopause, BMI, height, lifetime smoking exposure, CRP and genetic liabilities to twin births and PCOS on ovarian cancer risk. There was little evidence to support causal effects of genetic liability to type 2 diabetes, parity, or circulating levels of 25-hydroxyvitamin D or sex hormone-binding globulin on overall or histotype-specific EOC.

Though historically considered a homogeneous disease with a single cellular origin, epithelial ovarian cancer is now recognized as heterogeneous, consisting of multiple histological subtypes each with its own distinct origins, morphological characteristics, and molecular alterations ^18,43-46^. The largely histotype-specific findings in this analysis using genetic variants as proxies to minimize confounding and avoid reverse causation bias thus help to extend these insights further by supporting distinct causal pathways across EOC histotypes.

Some of the histotype-specific findings are consistent with conventional observational studies. For example, in agreement with previous analyses ^7-10,47-49^, most risk factors did not show clear evidence of association with HGSC. Consistent with some studies, age at natural menopause was most strongly associated with endometrioid carcinoma ^8^ and height was most strongly associated with clear cell carcinoma ^50,51^. The effect of genetic liability to endometriosis on risk of epithelial ovarian cancer is in agreement with two large pooled observational analyses ^9,52^, though these studies also reported positive risk relationships with endometrioid and low grade serous carcinoma.

However, some MR estimates were not consistent with those observed in conventional analyses. Most notably, previously reported associations between smoking and mucinous carcinoma ^9,53-55^ were not corroborated in MR analyses of lifetime smoking exposure. Though estimates from primary and sensitivity analyses all included the null line, inconsistencies in effect estimates across these analyses support pleiotropic biases distorting the causal effect estimate. Though parity has been consistently inversely associated with risk of ovarian cancer in conventional analyses ^10,56-60^, MR effect estimates suggesting a protective effect of giving birth to more children were imprecise and 95% confidence intervals spanned the null line. Given the few SNPs available to proxy for parity (two independent variants in this analysis), these results likely reflect limited statistical power.

Weaker statistical evidence also suggested an unexpected inverse effect of CRP, a marker of systemic inflammation, on endometrioid carcinoma and positive risk relationships between genetic liability to twin births and clear cell carcinoma. Given recent evidence to suggest a role of infectious agents in ovarian cancer [66, 67], a possible protective effect of CRP on endometrioid carcinoma could speculatively reflect the involvement of CRP in acute immune response (i.e., protection against active bacterial and viral infections). Meanwhile, the effect of genetic liability to twin births on clear cell carcinoma could be mediated by the higher levels of gonadotropins in the fertile years of women with a history of multiple births [54-56].

Overall, few previously reported risk factors showed clear evidence of a causal role in EOC or high grade serous carcinoma, the most common (∽70% of cases) and lethal EOC histotype, suggesting that some previously reported associations may have been driven by residual confounding, misclassification biases, or reverse causation ^61^. A notable exception was suggestive evidence that smoking increased odds of HGSC, consistent with some ^62,63^, but not all ^9,53,64,65^, observational analyses. A causal effect of genetic liability to endometriosis on EOC corroborates findings from conventional analyses that women with this condition are at elevated risk of subsequent disease^9,66^. This finding also suggests that subclinical manifestations of endometriosis may influence oncogenesis, indicating important avenues for future mechanistic work.

Strengths of this analysis include the use of a systematic approach to collate previously reported risk factors for EOC, the appraisal of the causal role of these risk factors in EOC etiology using a Mendelian randomization framework to reduce confounding and avoid reverse causation bias, the employment of complementary sensitivity analyses to rigorously assess for violations of MR assumptions, and the restriction of datasets utilized to women of primarily or exclusively European descent to minimize confounding through population stratification.

There are several limitations to these analyses. First, though F-statistics generated for most risk factors suggested that results were unlikely to suffer from weak instrument bias, statistical power for some analyses of less common ovarian cancer subtypes (low grade serous, mucinous, and clear cell carcinomas) was likely modest, meaning that the possibility that some results may reflect “false negative” findings cannot be ruled out. Since analyses were performed using summarized genetic association data in aggregate, it was not possible to restrict age at natural menopause analyses exclusively to participants who had undergone menopause. However, given that most ovarian cancer cases occur after menopause and that age-matched controls were used, the inclusion of some pre-or perimenopausal women in these analyses would likely have biased results toward the null (i.e., providing a conservative effect estimate). Additionally, models employed assumed no interaction (e.g., gene-environment, gene-gene) or effect modification and linear relationships between risk factors and ovarian cancer. Lastly, the use of a MR framework precluded directly examining the causal effects of some ovarian cancer risk factors that do not have robust genetic variants available to serve as proxies (e.g., use of oral contraceptives, hormone replacement therapy).

Though the largely null findings for overall EOC in this analysis can assist in de-prioritizing certain intervention targets for ovarian cancer prevention, they also underscore the challenges in establishing effective primary prevention strategies for this malignancy. To date, beyond risk-reducing surgical interventions, only the oral contraceptive pill has shown compelling evidence that regular use can reduce risk of subsequent disease ^59,67,68^. The continued identification of robust genetic variants to proxy other lifestyle and molecular factors previously reported to influence ovarian cancer (e.g., additional sex hormones, gonadotropins, inflammatory markers) will allow for a more refined assessment of the causal influence of these factors in ovarian carcinogenesis ^48,69^. Additionally, further work understanding possible mechanisms through which factors that appear to causally influence ovarian cancer in these analyses promote oncogenesis (e.g., genetic liability to endometriosis, C-reactive protein levels) could help to increase scope for prevention opportunities across the life-course. Lastly, for the vast majority of women who develop ovarian cancer with no previous history of smoking and who do not have endometriosis ^9,53,70^, there is a need to identify novel modifiable risk factors for this condition, as has been advocated elsewhere ^71,72^.

## Conclusions

Of 13 previously reported risk factors examined for association with overall epithelial ovarian cancer, only genetic liability to endometriosis and lifetime smoking exposure showed evidence compatible with a causal effect on disease risk. When stratified on ovarian cancer histotype, most risk factors showed causal effects on one or more subtypes, underscoring the heterogeneous nature of this disease. While this etiological heterogeneity could have implications for understanding mechanisms of tumour pathology and for studies examining histotype-specific prognosis, given the low incidence of EOC in the general population, prevention strategies targeting factors causally implicated in overall EOC are most likely to confer important population-level reductions in disease incidence. Along with effective clinical management of endometriosis and policies to prevent the initiation of tobacco use and encourage smoking cessation, established prevention strategies like the use of oral contraceptives continue to be important EOC risk-reducing mechanism. The identification of novel modifiable risk factors remains an important priority for the control of epithelial ovarian cancer.

## Acknowledgments

The authors’ responsibilities were as follows—JY, CLR, RMM, and SJL: conceived the study; JY: planned the analyses, conducted data analysis, and prepared the manuscript; JY, CLR, AL, KM, UM, AG, AW, JZ, GDS, RMM, SJL: critically revised the manuscript; and all authors: read and approved the final manuscript. None of the authors had any conflicts of interest to declare. The authors would like to thank the participants of the individual studies contributing to the Ovarian Cancer Association Consortium (OCAC) for their participation in these studies along with the principal investigators of OCAC for generating the data utilized for this analysis and for making this data available in the public domain.

## Footnote to Supplementary Figure 1

GWAS = genome-wide association study, SNP = single-nucleotide polymorphism, MR = Mendelian randomization, BMI = body mass index, CRP = C-reactive protein, SHBG = sex hormone-binding globulin

## Footnote to Figures 1-5

BMI = body mass index, PCOS = polycystic ovary syndrome, 25(OH)D = 25-hydroxyvitamin D, CRP = C-reactive protein, SHBG = sex hormone-binding globulin

