## Extended Methods for "Evaluating causal associations between previously reported risk factors and epithelial ovarian cancer: a Mendelian randomization analysis"

**GWAS and genetic instrument selection:**

If multiple genome-wide association studies (GWAS) were available for a previously reported risk factor, data was obtained from the largest GWAS available (defined as the total number of participants for continuous risk factors and total numbers of cases for binary risk factors). For risk factors derived from GWAS consisting of discovery and replication analyses, effect estimates were obtained from combined discovery-replication GWAS meta-analyses. For age at menarche and twin pregnancy analyses, as effect estimates were presented separately for discovery and replication analyses (but not for the pooled analysis), effect estimates were obtained from the replication phase of the analysis to minimize “winner’s curse” bias (exaggerated effect sizes for SNPs identified in the discovery stage of a GWAS due to chance correlation between genetic variants and confounders).

After obtaining effect estimates from relevant GWAS, for risk factors with two or more SNPs available, SNPs were pruned for linkage disequilibrium at r^2^ < 0.001 at a clumping distance of 10,000 kilobases from the lead SNP at *P* < 5 x 10^-8^ with reference to the 1000 Genomes Project (<http://www.internationalgenome.org/>). Of the remaining independent SNPs, corresponding effect estimates and standard errors for these SNPs were then obtained from ovarian cancer datasets. When a SNP was not available in the ovarian cancer dataset, the presence of a “proxy” SNP in linkage disequilibrium with this SNP at r^2^≥0.80 was assessed using the 1000 genomes European sample data. If such “proxy” SNP was not available in the ovarian cancer dataset, this SNP was removed from the analysis.

**Sensitivity analyses to examine horizontal pleiotropy:**

The following sensitivity analyses were performed to examine horizontal pleiotropy in Mendelian randomization (MR) analyses: MR-Egger, weighted median estimator, and weighted mode estimator. Descriptions of these sensitivity analyses along with their assumptions are provided below:

*MR-Egger:*

MR-Egger relaxes the exclusion restriction criterion and thus can provide unbiased estimates of causal effects even when all single-nucleotide polymorphisms (SNPs) in an instrument are invalid through violation of this assumption. This approach performs a weighted generalized linear regression of the SNP-outcome effect estimates on the SNP-exposure effect estimates with an unconstrained intercept term (i.e., unconstrained to pass through zero). Provided that the InSIDE (Instrument Strength Independent of Direct Effect) assumption is met (that no association exists between the strength of SNP-risk factor associations and the strength of bias due to horizontal pleiotropy) and that measurement error in the genetic instrument is negligible (“No Measurement Error” or NOME assumption), the slope generated from MR-Egger regression can provide an estimate of the causal effect of a risk factor on a disease outcomes that is adjusted for directional pleiotropy (where the horizontally pleiotropic effect across a genetic instrument do not average to zero) and the intercept term can provide a formal statistical test for directional pleiotropy.

*Weighted median estimator*

The weighted median estimator (WME) approach provides an estimate of the weighted median of a distribution in which individual SNP effect estimates in an instrument are ordered and weighted by the inverse of their variance. Unlike MR-Egger which can provide an unbiased causal effect even when all SNPs are invalid instruments, WME requires that at least 50% of the information in a multi-allelic instrument is coming from SNPs that are valid instrumental variables in order to provide an unbiased estimate of a causal effect in an MR analysis. However, the WME has two advantages over MR-Egger in that it provides improved precision as compared to the latter and does not rely on the InSIDE assumption.

*Weighted mode estimator*

The weighted mode-based estimator generates a causal estimate using the mode of a smoothed empirical density function of individual SNP effect estimates in a multi-allelic instrument, weighted by the inverse variance of the SNP-outcome association. This approach operates under the assumption that the most common effect estimate of individual SNPs in a multi-allelic instrument arises from valid instruments (called the Zero Modal Pleiotropy Assumption, or ZEMPA). If this assumption holds, the mode can provide a consistent causal estimate even if most of the (non-modal) SNPs are invalid. Mode-based approaches have less power to detect a causal effect than the weighted median estimator but greater power than MR-Egger regression under the condition of no invalid instruments. Similar to the weighted median estimator, mode-based approaches are also (by default) less susceptible to bias from outlying variants in a risk score.

**Assessment of sex-specific instruments:**

For GWAS of body mass index, height, type 2 diabetes, parity, 25-hydroxyvitamin D, l-ascorbic acid, lifetime smoking exposure, C-reactive protein, and sex hormone-binding globulin, evidence for heterogeneity of SNP-risk factor associations by sex was examined. For GWAS that performed a statistical test for interaction by sex for SNPs that achieved genome-wide significance, the reported *P*-value for interaction by sex was compared to a Bonferroni-corrected *P*-value that adjusted for multiple assessment tests for interaction. For example, tests for interaction by sex were performed for 8 genome-wide significant SNPs identified in the sex hormone-binding globulin (SHBG) GWAS. After correcting for “look up” of 8 statistical tests (0.05/8=0.0063), 0 of 8 genome-wide significant SNPs associated with SHBG showed evidence of interaction by sex. SNP-SHBG effect estimates were correspondingly obtained from from sex-combined analyses. Likewise, after multiple look-up correction (0.05/9=0.0063), 0 of 9 genome-wide significant SNPs associated with c-reactive protein (CRP) showed evidence of interaction by sex. As only 2 of 66 BMI SNPs (0.05/66=0.00076) showed evidence for interaction by sex after correcting for multiple look-up, it was considered unlikely that MR estimates would be biased by using effect estimates from sex-combined analyses for these 2 SNPs and, therefore, sex-combined effect estimates were used for all SNPs. 0 of 10 (0.05/10=0.005) type 2 diabetes SNPs showed evidence for sex-specific effects and, thus, sex-combined effect estimates in genetic liability to type 2 diabetes were used for MR analyses. Likewise, 0 of 2 (0.05/2= 0.025) parity SNPs showed evidence for sex-specific effects and, therefore, sex-combined effect estimates were used for MR analyses of parity. Interaction by sex was not assessed in genome-wide association studies for height, lifetime smoking exposure, or 25-hydroxyvitamin D.

**Risk-factor specific sensitivity analyses:**

As an additional sensitivity analysis for C-reactive protein (CRP) analyses, MR analyses were performed using a conservative genetic instrument that consisted exclusively of SNPs located in the *CRP* gene region. In total, 4 SNPs (rs1130864, rs1205, rs1800947, rs3093077) were used that were in mild linkage disequilibrium with each other (r^2^≤0.20), using the 1000 Genomes Project Phase 3 as reference. Compared to the primary instrument for CRP (8 SNPs robustly associated with CRP from different gene regions), SNPs identified within *CRP* are more likely to influence CRP levels directly and thus may be less susceptible to horizontal pleiotropic bias.
