## Supplementary figures and images for "Evaluating causal associations between previously reported risk factors and epithelial ovarian cancer: a Mendelian randomization analysis"

### Supplementary Figure 1

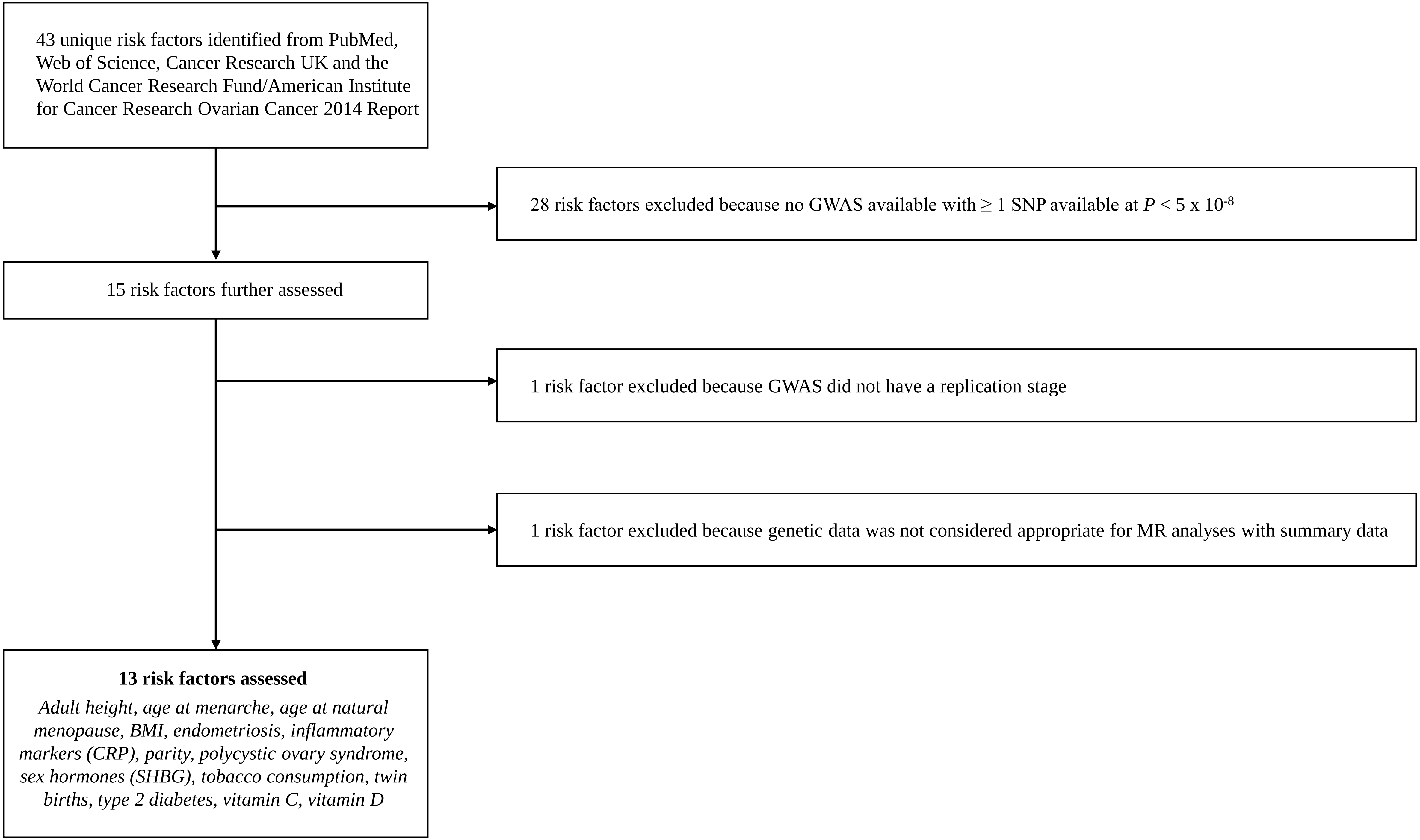
