## Supplementary Materials for "Evaluating causal associations between previously reported risk factors and epithelial ovarian cancer: a Mendelian randomization analysis"

**Ovarian Cancer Association Consortium (OCAC) study population and OncoArray genome-wide association analyses:**

The OCAC OncoArray data comprised 63 genotyping project/case-controls sets. Some studies contributed samples to more than one genotyping project and some case-control sets are a combination of multiple individual studies. Genotype data were obtained by either direct genotyping using an Illumina Custom Infinium array (OncoArray) consisting of approximately 530,000 SNPs or by imputation with reference to the 1000 Genomes Project Phase Three dataset. All SNPs with a call rate of <95%, evidence of violation of Hardy-Weinberg equilibrium (*P*<10^-7^ in controls or *P*<10^-12^ in cases), or SNPs with a concordance<98% among 5,280 duplicate pairs were removed. In imputation, all SNPs with a minor allele frequency of <1% and a call rate of <98% and SNPs that could not be linked to the 1000 genomes reference or different significantly in frequency from the 1000 genomes (European frequency) in addition to 1,128 SNPs where the cluster plot was judged to be inadequate were removed. Risk analyses for ovarian cancer outcomes were performed across OncoArray analyses in addition to COGS (a previous GWAS meta-analysis of ovarian cancer) and five GWAS datasets using logistic regression in models that were adjusted for study and population substructure by including eigenvectors of project-specific principal components of ancestry as covariates in the model. Summary estimates were meta-analyzed across studies using an inverse-variance fixed-effects approach.

**Search strategies for PubMed and Web of Science:**

PubMed:

((ovarian cancer[MeSH Terms]) AND epidemiology) AND review[Filter] Filters: Review; Publication date from 2012/02/13 to 2017/02/13

Web of Science:

((TS=("ovarian cancer") AND TS=(epidemiology))) *AND* **LANGUAGE:** (English) *AND* **DOCUMENT TYPES:** (Review)

*Indexes=SCI-EXPANDED Timespan=2012-2017*

(Searched on 02/10/2017)

**Review papers identified from PubMed and Web of Science searches:**

Bowtell DD, Böhm S, Ahmed AA, et al. Rethinking ovarian cancer II: reducing mortality from high-grade serous ovarian cancer. Nat Rev Cancer. 2015 Nov;15(11):668-79.

Crane TE, Khulpateea BR, Alberts DS, Basen-Engquist K, Thomson CA. Dietary intake and ovarian cancer risk: a systematic review. Cancer Epidemiol Biomarkers Prev. 2014 Feb;23(2):255-73.

Hunn J, Rodriguez GC. Ovarian cancer: etiology, risk factors, and epidemiology. Clin Obstet Gynecol. 2012 Mar;55(1):3-23.

Karnezis AN, Cho KR, Gilks CB, Pearce CL, Huntsman DG. The disparate origins of ovarian cancers: pathogenesis and prevention strategies. Nat Rev Cancer. 2017 Jan;17(1):65-74.

Tropé CG, Kaern J, Davidson B. Borderline ovarian tumours. Best Pract Res Clin Obstet Gynaecol. 2012 Jun;26(3):325-36.

Webb PM, Jordan SJ. Epidemiology of epithelial ovarian cancer. Best Pract Res Clin Obstet Gynaecol. 2017 May;41:3-14. doi: 10.1016/j.bpobgyn.

**Risk factor inclusion steps:**

43 unique risk factors identified from PubMed, Web of Science, Cancer Research UK and the World Cancer Research Fund/American Institute for Cancer Research Ovarian Cancer 2014 Report

Adult height, age at first child birth, age at last birth, age at menarche, age at natural menopause, asbestos exposure, aspirin intake, body mass index, breastfeeding, dairy intake, dietary fat, endometriosis, family history of ovarian cancer, fruit intake, gamma radiation, hormone replacement therapy (oestrogen only), hormone replacement therapy (oestrogen-progestogen), hysterectomy, infertility, inflammatory markers, isoflavone intake, lactose intake, metformin, nitrites, oestrogen-progestogen contraceptives, opportunistic salpingectomy, parity, pelvic inflammatory disease, physical activity, polycystic ovary syndrome, risk-reducing salpingo-oophorectomy, sedentary behaviour, sex hormones, talc-based body powder, tea consumption, tobacco consumption, tubal ligation, twin births, type 2 diabetes, vegetable intake, vitamin C, vitamin D, x-radiation

28 risk factors excluded because no GWAS available with ≥ 1 SNP at *P* < 5 x 10^-8^

Age at last birth, asbestos exposure, aspirin intake, breastfeeding, dairy intake, dietary fat, family history of ovarian cancer, fruit intake, gamma radiation, hormone replacement therapy (oestrogen only), hormone replacement therapy (oestrogen-progestogen), hysterectomy, infertility, isoflavone intake, lactose intake, metformin, nitrites, oestrogen-progestogen contraceptives, opportunistic salpingectomy, pelvic inflammatory disease, risk-reducing salpingo-oophorectomy, sedentary behaviour, talc-based body powder, tea consumption, tubal ligation, vegetable intake, vitamin C, x-radiation

1 risk factor excluded because GWAS identified did not have replication stage or was not a meta-analysis

Physical activity (https://doi.org/10.1101/261719)

1 risk factor excluded because genetic data was not considered appropriate for Mendelian randomization analyses with summary data

Age at first child birth (https://doi.org/10.1038/ng.3698)

13 risk factors assessed

Adult height, age at menarche, age at natural menopause, body mass index, endometriosis, inflammatory markers (C-reactive protein), parity, polycystic ovary syndrome, sex hormones (sex hormone-binding globulin), tobacco consumption, twin births, type 2 diabetes, vitamin D
