## Supplementary Table 1 for "Evaluating causal associations between previously reported risk factors and epithelial ovarian cancer: a Mendelian randomization analysis"

| **Reported risk factor** | **GWAS** | **Sex** | **No. of SNPs**  **available^1^** | **No. of SNPs used^2^** | **R^2^** | **F-statistic** |
| --- | --- | --- | --- | --- | --- | --- |
| **Reproductive factors** | |  |  |  |  |  |
| Age at menarche | Day et al (2017)^1^ | Women | 368 | 317 | 5.1 | 55 |
| Age at natural menopause | Day et al (2015)^2^ | Women | 42 | 35 | 4.2 | 86 |
| Twin births | Mbarek et al (2016)^3^ | Women | 2 | 2 | 0.06 | 4 |
| Parity | Barban et al (2016)^4^ | Combined | 2 | 2 | 0.02 | 33 |
| **Anthropometric traits** | |  |  |  |  |  |
| Body mass index | Locke et al. (2015)^5^ | Combined | 77 | 66 | 2.2 | 110 |
| Height | Wood et al. (2014)^6^ | Combined | 386 | 345 | 11.8 | 98 |
| **Clinical factors** | |  |  |  |  |  |
| Type 2 diabetes | Morris et al. (2012)^7^ | Combined | 10 | 10 | 0.34 | 24 |
| Endometriosis | Sapkota et al (2017)^8^ | Women | 10 | 10 | 0.24 | 49 |
| Polycystic ovary syndrome | Day et al. (2018)^9^ | Women | 14 | 11 | 0.35 | 36 |
| **Lifestyle factors** | |  |  |  |  |  |
| 25-hydroxyvitamin D | Jiang et al (2018)^10^ | Combined | 6 | 5 | 2.6 | 423 |
| Lifetime smoking exposure | Wooton et al (2018)^11^ | Combined | 124 | 115 | 1.2 | 49 |
| **Molecular risk factors** |  |  |  |  |  |  |
| C-reactive protein | Deghan et al (2011)^12^ | Combined | 9 | 8 | 3.6 | 382 |
| SHBG | Coviello et al. (2012)^13^ | Combined | 9 | 8 | 4.3 | 121 |

**Supplementary Table 1. Catalogue of GWAS used for genetic instruments and estimates of instrument strength for previously reported risk factors**

^1^ Corresponds to the number of SNPs available after pruning top SNPs reported in corresponding linkage disequilibrium at r^2^ < 0.001 at a clumping distance of 10,000 kilobases from the lead SNP at *P* < 5 x 10^-8^

^2^ Corresponds to the number of SNPs (or linkage disequilibrium proxies) available in ovarian cancer datasets

GWAS = genome-wide association study, SNPs = single-nucleotide polymorphisms, SHBG = sex hormone-binding globulin.

R^2^ represents the proportion of variance in a risk factor explained by the genetic instrument. F-statistic represents the strength of the association between the genetic instrument and levels of the risk factor.
