## Supplementary Tables 2-6 for "Evaluating causal associations between previously reported risk factors and epithelial ovarian cancer: a Mendelian randomization analysis"

| **Risk factor** | **Ovarian cancer outcome** | **IVW**  **OR (95% CI)** | ***P*-value** | **MR-Egger regression**  **OR (95% CI)** | ***P*-value** | **MR-Egger intercept**  **OR (95% CI)** | ***P*-value** | **Weighted median**  **OR (95%CI)** | ***P*-value** | **Weighted mode**  **OR (95% CI)** | ***P*-value** |
| --- | --- | --- | --- | --- | --- | --- | --- | --- | --- | --- | --- |
| **Age at menarche** | | | | | | | | | | | |
|  | Overall | 1.07 (1.00-1.14) | 0.046 | 1.00 (0.89-1.13) | 0.94 | 1.00 (1.00-1.01) | 0.22 | 1.01 (0.92-1.10) | 0.88 | 0.98 (0.25-3.84) | 0.98 |
|  | HGSC | 1.05 (0.97-1.13) | 0.24 | 0.94 (0.82-1.08) | 0.36 | 1.00 (1.00-1.01) | 0.06 | 0.99 (0.88-1.11) | 0.82 | 1.03 (0.29-3.63) | 0.97 |
|  | LGSC | 1.08 (0.86-1.36) | 0.50 | 0.93 (0.61-1.41) | 0.73 | 1.01 (0.99-1.02) | 0.39 | 1.03 (0.72-1.46) | 0.88 | 1.33 (2.5e^-5^-7.2e^4^) | 0.38 |
|  | Mucinous | 1.13 (0.95-1.35) | 0.18 | 1.31 (0.95-1.81) | 0.10 | 0.99 (0.98-1.00) | 0.29 | 1.18 (0.89-1.56) | 0.26 | 0.74 (9.5e^-6^-5.9e^4^) | 0.96 |
|  | Endometrioid | 1.19 (1.05-1.36) | 0.008 | 1.17 (0.93-1.48) | 0.19 | 1.00 (0.99-1.01) | 0.86 | 1.15 (0.94-1.41) | 0.17 | 1.26 (0.12-12.8) | 0.85 |
|  | Clear cell | 1.04 (0.87-1.25) | 0.66 | 0.91 (0.71-1.37) | 0.94 | 1.00 (0.99-1.01) | 0.70 | 0.94 (0.69-1.27) | 0.68 | 0.93 (0.02-35.6) | 0.97 |
|  | LMP | 1.06 (0.94-1.21) | 0.35 | 0.93 (0.74-1.17) | 0.55 | 1.00 (1.00-1.01) | 0.18 | 1.05 (0.86-1.28) | 0.66 | 1.02 (0.09-12.0) | 0.99 |
| **Age at natural menopause** | | | | | | | | | | | |
|  | Overall | 1.03 (1.00-1.06) | 0.07 | 1.07 (1.00-1.14) | 0.06 | 0.99 (0.98-1.00) | 0.20 | 1.03 (0.99-1.07) | 0.10 | 1.04 (1.00-1.09) | 0.06 |
|  | HGSC | 1.02 (0.99-1.05) | 0.31 | 1.03 (0.97-1.11) | 0.35 | 1.00 (0.98-1.01) | 0.58 | 1.02 (0.98-1.07) | 0.39 | 1.02 (0.97-1.08) | 0.36 |
|  | LGSC | 1.01 (0.92-1.12) | 0.77 | 1.10 (0.89-1.36) | 0.36 | 0.98 (0.94-1.02) | 0.39 | 1.08 (0.95-1.23) | 0.26 | 1.07 (0.92-1.26) | 0.38 |
|  | Mucinous | 1.00 (0.92-1.09) | 0.97 | 1.00 (0.82-1.23) | 0.99 | 1.00 (0.96-1.04) | 0.98 | 1.03 (0.92-1.15) | 0.58 | 1.02 (0.90-1.15) | 0.81 |
|  | Endometrioid | 1.09 (1.02-1.16) | 0.007 | 1.13 (0.98-1.30) | 0.10 | 0.99 (0.97-1.02) | 0.57 | 1.11 (1.02-1.20) | 0.01 | 1.11 (1.02-1.21) | 0.03 |
|  | Clear cell | 1.05 (0.96-1.14) | 0.29 | 1.26 (1.05-1.52) | 0.02 | 1.00 (0.96-1.04) | 0.03 | 1.11 (0.99-1.25) | 0.08 | 1.16 (1.02-1.31) | 0.03 |
|  | LMP | 1.04 (0.98-1.10) | 0.21 | 1.11 (0.98-1.25) | 0.12 | 0.99 (0.96-1.01) | 0.26 | 1.08 (0.99-1.17) | 0.07 | 1.10 (1.00-1.20) | 0.05 |
| **Genetic liability to twin births** | | | | | | | | | | | |
|  | Overall | 1.10 (0.85-1.43) | 0.45 | - | - | - | - | - | - | - | - |
|  | HGSC | 1.03 (0.78-1.37) | 0.81 | - | - | - | - | - | - | - | - |
|  | LGSC | 1.23 (0.65-2.30) | 0.53 | - | - | - | - | - | - | - | - |
|  | Mucinous | 0.72 (0.43-1.22) | 0.22 | - | - | - | - | - | - | - | - |
|  | Endometrioid | 1.88 (1.00-3.53) | 0.05 | - | - | - | - | - | - | - | - |
|  | Clear cell | 1.78 (1.05-3.03) | 0.03 | - | - | - | - | - | - | - | - |
|  | LMP | 1.32 (0.91-1.92) | 0.15 |  |  |  |  |  |  |  |  |
| **Parity** |  |  |  |  |  |  |  |  |  |  |  |
|  | Overall | 0.66 (0.26-1.69) | 0.39 | - | - | - | - | - | - | - | - |
|  | HGSC | 0.90 (0.30-2.72) | 0.85 | - | - | - | - | - | - | - | - |
|  | LGSC | 0.74 (0.03-21.7) | 0.86 | - | - | - | - | - | - | - | - |
|  | Mucinous | 7.65(0.47-124.8) | 0.15 | - | - | - | - | - | - | - | - |
|  | Endometrioid | 0.19 (0.01-3.23) | 0.25 | - | - | - | - | - | - | - | - |
|  | Clear cell | 0.15 (0.02-1.11) | 0.06 | - | - | - | - | - | - | - | - |
|  | LMP | 1.89 (0.26-14.0) | 0.53 |  |  |  |  |  |  |  |  |

**Supplementary Table 2. IVW and sensitivity analysis estimates for causal estimates of reproductive traits on ovarian cancer risk**

Causal estimates are scaled to represent the effect of a one-year decrease in age at onset for menarche, a one-year increase in age at onset for natural menopause, a one-unit higher log odds liability to having spontaneous dyzgotic twins, and a one-child increase in number of children ever born. IVW = Inverse-variance weighted, HGSC = High grade serous carcinoma, LGSC = Low grade serous carcinoma, LMP = Low malignant potential.

**Supplementary Table 3. IVW and sensitivity analysis estimates for causal estimates of anthropometric traits on ovarian cancer risk**

| **Risk factor** | **Ovarian cancer outcome** | **IVW**  **OR (95% CI)** | ***P*-value** | **MR-Egger regression**  **OR (95% CI)** | ***P*-value** | **MR-Egger intercept**  **OR (95% CI)** | ***P*-value** | **Weighted median**  **OR (95%CI)** | ***P*-value** | **Weighted mode**  **OR (95% CI)** | ***P*-value** |
| --- | --- | --- | --- | --- | --- | --- | --- | --- | --- | --- | --- |
| **Body mass index** | | | | | | | | | | | |
|  | Overall | 1.23 (1.07-1.42) | 0.003 | 1.32 (0.88-1.99) | 0.19 | 1.00 (0.99-1.01) | 0.73 | 1.14 (0.93-1.40) | 0.21 | 1.06 (0.75-1.51) | 0.18 |
|  | HGSC | 1.26 (1.06-1.50) | 0.01 | 1.05 (0.63-1.75) | 0.85 | 1.01 (0.99-1.02) | 0.47 | 1.17 (0.91-1.50) | 0.21 | 0.95 (0.53-1.35) | 0.49 |
|  | LGSC | 1.27 (0.73-2.20) | 0.40 | 3.35 (0.68-16.6) | 0.14 | 0.97 (0.93-1.02) | 0.21 | 1.51 (0.71-3.22) | 0.28 | 1.46 (0.45-4.79) | 0.53 |
|  | Mucinous | 1.15 (0.73-1.79) | 0.55 | 4.06 (1.15-14.4) | 0.03 | 0.96 (0.93-1.00) | 0.04 | 1.21 (0.65-2.24) | 0.55 | 1.23 (0.44-3.48) | 0.69 |
|  | Endometrioid | 1.48 (1.07-2.06) | 0.02 | 2.31 (0.89-6.00) | 0.09 | 0.99 (0.96-1.01) | 0.34 | 1.98 (1.23-3.19) | 0.005 | 2.38 (0.86-6.61) | 0.10 |
|  | Clear cell | 0.83 (0.55-1.26) | 0.39 | 1.41 (0.42-4.65) | 0.58 | 0.99 (0.95-1.02) | 0.37 | 1.36 (0.74-2.50) | 0.33 | 1.82 (0.58-5.70) | 0.31 |
|  | LMP | 1.39 (1.04-1.85) | 0.03 | 1.38 (0.60-3.20) | 0.45 | 1.00 (0.98-1.02) | 0.99 | 1.29 (0.83-1.99) | 0.25 | 1.17 (0.55-2.49) | 0.68 |
| **Height** | | | | | | | | | | | |
|  | Overall | 1.02 (0.96-1.08) | 0.47 | 1.10 (0.95-1.29) | 0.21 | 1.00 (0.99-1.00) | 0.29 | 1.00 (0.92-1.10) | 0.92 | 0.99 (0.84-1.18) | 0.92 |
|  | HGSC | 1.00 (0.94-1.08) | 0.93 | 1.11 (0.92-1.35) | 0.26 | 1.00 (0.99-1.00) | 0.24 | 0.93 (0.84-1.03) | 0.17 | 0.91 (0.74-1.14) | 0.42 |
|  | LGSC | 0.90 (0.74-1.09) | 0.27 | 1.06 (0.63-1.79) | 0.83 | 0.99 (0.98-1.01) | 0.50 | 0.87 (0.64-1.17) | 0.35 | 0.74 (0.40-1.37) | 0.33 |
|  | Mucinous | 0.99 (0.85-1.17) | 0.94 | 1.10 (0.71-1.69) | 0.67 | 1.00 (0.98-1.01) | 0.63 | 1.03 (0.81-1.31) | 0.79 | 1.12 (0.69-1.83) | 0.65 |
|  | Endometrioid | 1.01 (0.90-1.14) | 0.87 | 0.88 (0.64-1.22) | 0.45 | 1.00 (0.99-1.01) | 0.38 | 1.01 (0.83-1.22) | 0.95 | 1.01 (0.66-1.55) | 0.96 |
|  | Clear cell | 1.36 (1.15-1.61) | 0.0003 | 1.20 (0.76-1.88) | 0.44 | 1.00 (0.99-1.02) | 0.55 | 1.38 (1.07-1.79) | 0.01 | 1.31 (0.72-2.41) | 0.38 |
|  | LMP | 1.09 (0.97-1.22) | 0.15 | 1.01 (1.00-1.02) | 0.20 | 0.90 (0.66-1.22) | 0.50 | 1.09 (0.92-1.30) | 0.32 | 1.27 (0.85-1.89) | 0.24 |

Causal estimates are scaled to represent the effect of a 1-SD increase in body mass index (kg/m^2^) and a 1-SD increase in height (cm). IVW = Inverse-variance weighted, HGSC = High grade serous carcinoma, LGSC = Low grade serous carcinoma, LMP = Low malignant potential.

**Supplementary Table 4. IVW and sensitivity analysis estimates for causal estimates of clinical factors on ovarian cancer risk**

| **Risk factor** | **Ovarian cancer outcome** | **IVW**  **OR (95% CI)** | ***P*-value** | **MR-Egger regression**  **OR (95% CI)** | ***P*-value** | **MR-Egger intercept**  **OR (95% CI)** | ***P*-value** | **Weighted median**  **OR (95%C)** | ***P*-value** | **Weighted mode**  **OR (95% CI)** | ***P*-value** |
| --- | --- | --- | --- | --- | --- | --- | --- | --- | --- | --- | --- |
| **Genetic liability to endometriosis** | | | | | | | | | | | |
|  | Overall | 1.27 (1.16-1.40) | 6.9e^-7^ | 1.73 (1.05-2.85) | 0.06 | 0.97 (0.92-1.02) | 0.25 | 1.16 (1.09-1.23) | 6.9e^-6^ | 1.17 (0.24-5.60) | 0.85 |
|  | HGSC | 1.17 (1.04-1.31) | 0.007 | 1.76 (0.97-3.17) | 0.10 | 0.96 (0.90-1.02) | 0.21 | 1.05 (0.83-1.33) | 0.70 | 0.98 (0.49-1.95) | 0.95 |
|  | LGSC | 1.26 (0.87-1.81) | 0.22 | 2.02 (0.27-14.9) | 0.51 | 0.95 (0.77-1.17) | 0.65 | 1.00 (0.37-2.68) | 0.99 | 1.00 (0.40-2.48) | 0.99 |
|  | Mucinous | 1.32 (0.94-1.85) | 0.11 | 1.12 (0.17-7.49) | 0.91 | 1.02 (0.84-1.24) | 0.87 | 1.03 (0.82-1.29) | 0.78 | 1.02 (0.46-2.23) | 0.96 |
|  | Endometrioid | 1.37 (1.10-1.69) | 0.004 | 0.89 (0.28-2.82) | 0.84 | 1.05 (0.93-1.18) | 0.48 | 1.03 (0.88-1.19) | 0.73 | 0.88 (0.11-6.74) | 0.90 |
|  | Clear cell | 2.69 (1.88-3.86) | 7.4e^-8^ | 4.37 (0.60-31.9) | 0.18 | 0.95 (0.77-1.17) | 0.64 | 1.56 (1.01-2.41) | 0.05 | 1.59 (0.02-131.9) | 0.84 |
|  | LMP | 1.33 (1.09-1.63) | 0.006 | 1.13 (0.39-3.28) | 0.83 | 1.02 (0.91-1.14) | 0.76 | 1.23 (1.07-1.41) | 0.003 | 1.23 (0.15-10.2) | 0.85 |
| **Genetic liability to polycystic ovary syndrome** | | | | | | | | | | | |
|  | Overall | 0.97 (0.87-1.09) | 0.64 | 0.83 (0.48-1.45) | 0.53 | 0.97 (0.92-1.02) | 0.25 | 0.93 (0.83-1.04) | 0.19 | 0.93 (0.79-1.09) | 0.37 |
|  | HGSC | 0.97 (0.85-1.11) | 0.69 | 0.99 (0.51-1.92) | 0.97 | 0.96 (0.90-1.02) | 0.21 | 0.96 (0.84-1.09) | 0.53 | 0.97 (0.80-1.17) | 0.73 |
|  | LGSC | 1.33 (1.01-1.74) | 0.04 | 1.00 (0.27-3.67) | 0.99 | 0.95 (0.77-1.17) | 0.65 | 1.39 (0.97-1.98) | 0.07 | 1.33 (0.77-2.28) | 0.33 |
|  | Mucinous | 1.15 (0.92-1.44) | 0.22 | 0.59 (0.20-1.72) | 0.36 | 1.02 (0.84-1.24) | 0.87 | 1.15 (0.84-1.57) | 0.38 | 1.16 (0.65-2.08) | 0.62 |
|  | Endometrioid | 0.74 (0.62-0.90) | 0.002 | 0.52 (0.21-1.31) | 0.20 | 1.05 (0.93-1.18) | 0.48 | 0.70 (0.55-0.88) | 0.003 | 0.65 (0.43-0.98) | 0.07 |
|  | Clear cell | 1.03 (0.80-1.32) | 0.82 | 0.73 (0.21-2.55) | 0.64 | 0.95 (0.77-1.17) | 0.64 | 1.01 (0.73-1.38) | 0.97 | 0.84 (0.49-1.42) | 0.52 |
|  | LMP | 1.08 (0.92-1.26) | 0.37 | 1.02 (0.48-2.19) | 0.95 | 1.01 (0.91-1.11) | 0.90 | 1.06 (0.86-1.30) | 0.60 | 1.05 (0.75-1.47) | 0.77 |
| **Genetic liability to type 2 diabetes** | | | | | | | | | | | |
|  | Overall | 0.99 (0.93-1.06) | 0.81 | 0.86 (0.59-1.26) | 0.46 | 1.02 (0.97-1.08) | 0.48 | 1.02 (0.93-1.11) | 0.72 | 1.05 (0.91-1.22) | 0.52 |
|  | HGSC | 0.95 (0.86-1.06) | 0.36 | 0.95 (0.51-1.78) | 0.83 | 1.00 (0.92-1.09) | 0.99 | 0.94 (0.83-1.05) | 0.28 | 0.90 (0.73-1.12) | 0.38 |
|  | LGSC | 0.94 (0.74-1.20) | 0.63 | 0.82 (0.22-3.07) | 0.77 | 1.02 (0.85-1.22) | 0.83 | 0.92 (0.66-1.27) | 0.60 | 0.85 (0.51-1.40) | 0.54 |
|  | Mucinous | 0.96 (0.79-1.17) | 0.70 | 0.61 (0.20-1.83) | 0.40 | 1.07 (0.92-1.24) | 0.43 | 0.96 (0.74-1.23) | 0.74 | 0.95 (0.66-1.37) | 0.78 |
|  | Endometrioid | 1.13 (0.94-1.35) | 0.20 | 0.80 (0.28-2.29) | 0.69 | 1.05 (0.91-1.21) | 0.54 | 1.05 (0.86-1.28) | 0.65 | 1.03 (0.77-1.38) | 0.83 |
|  | Clear cell | 0.97 (0.78-1.19) | 0.75 | 0.58 (0.17-1.95) | 0.41 | 1.07 (0.91-1.27) | 0.43 | 1.01 (0.77-1.32) | 0.95 | 1.06 (0.64-1.74) | 0.83 |
|  | LMP | 1.14 (0.97-1.33) | 0.11 | 1.29 (0.51-3.29) | 0.61 | 0.98 (0.86-1.12) | 0.80 | 1.12 (0.92-1.36) | 0.25 | 1.12 (0.80-1.56) | 0.53 |

Causal estimates are scaled to represent the effect of a one-unit log odds higher liability to type 2 diabetes, endometriosis, or polycystic ovary syndrome. IVW = Inverse-variance weighted, HGSC = High grade serous carcinoma, LGSC = Low grade serous carcinoma, LMP = Low malignant potential.

**Supplementary Table 5. IVW and sensitivity analysis estimates for causal estimates of lifestyle factors on ovarian cancer risk**

| **Risk factor** | **Ovarian cancer outcome** | **IVW**  **OR (95% CI)** | ***P*-value** | **MR-Egger regression**  **OR (95% CI)** | ***P*-value** | **MR-Egger intercept**  **OR (95% CI)** | ***P*-value** | **Weighted median**  **OR (95%C)** | ***P*-value** | **Weighted mode**  **OR (95% CI)** | ***P*-value** |
| --- | --- | --- | --- | --- | --- | --- | --- | --- | --- | --- | --- |
| **25-hydroxyvitamin D** | | | | | | | | | | | |
|  | Overall | 1.02 (0.72-1.44) | 0.93 | 1.48 (0.82-2.69) | 0.28 | 0.98 (0.95-1.01) | 0.24 | 1.15 (0.85-1.57) | 0.36 | 1.16 (0.84-1.61) | 0.41 |
|  | HGSC | 0.98 (0.61-1.56) | 0.92 | 1.68 (0.79-3.59) | 0.27 | 0.97 (0.93-1.01) | 0.20 | 1.09 (0.77-1.54) | 0.63 | 1.14 (0.80-1.61) | 0.51 |
|  | LGSC | 0.64 (0.24-1.75) | 0.39 | 1.62 (0.24-11.1) | 0.66 | 0.95 (0.86-1.04) | 0.35 | 0.71 (0.24-2.11) | 0.54 | 0.76 (0.23-2.46) | 0.67 |
|  | Mucinous | 1.27 (0.56-2.87) | 0.57 | 0.87 (0.18-4.23) | 0.87 | 1.02 (0.95-1.11) | 0.62 | 1.19 (0.50-2.85) | 0.70 | 1.17 (0.45-2.99) | 0.76 |
|  | Endometrioid | 0.83 (0.46-1.50) | 0.54 | 0.99 (0.31-3.11) | 0.98 | 0.99 (0.94-1.05) | 0.76 | 0.92 (0.49-1.72) | 0.78 | 0.93 (0.45-1.89) | 0.84 |
|  | Clear cell | 1.75 (0.77-3.99) | 0.18 | 2.41 (0.49-11.8) | 0.36 | 0.98 (0.91-1.06) | 0.68 | 2.00 (0.85-4.72) | 0.11 | 1.97 (0.77-5.06) | 0.23 |
|  | LMP | 1.02 (0.43-2.42) | 0.96 | 0.39 (0.09-1.62) | 0.28 | 1.06 (0.99-1.14) | 0.22 | 0.85 (0.43-1.68) | 0.65 | 0.75 (0.39-1.47) | 0.45 |
| **Comprehensive smoking exposure** | | | | | | | | | | | |
|  | Overall | 1.36 (1.04-1.78) | 0.02 | 1.83 (0.65-5.17) | 0.26 | 1.00 (0.99-1.01) | 0.57 | 1.38 (0.93-2.06) | 0.11 | 1.40 (0.55-3.59) | 0.48 |
|  | HGSC | 1.44 (1.05-1.98) | 0.02 | 2.67 (0.78-9.16) | 0.12 | 0.99 (0.98-1.01) | 0.31 | 1.65 (1.02-2.67) | 0.04 | 2.15 (0.78-5.89) | 0.14 |
|  | LGSC | 1.04 (0.38-2.88) | 0.94 | 0.06 (0.00-2.90) | 0.16 | 1.03 (0.99-1.07) | 0.14 | 0.69 (0.17-2.87) | 0.61 | 0.19 (0.01-4.54) | 0.31 |
|  | Mucinous | 1.41 (0.63-3.17) | 0.41 | 0.50 (0.02-11.9) | 0.67 | 1.01 (0.98-1.05) | 0.51 | 0.92 (0.27-3.11) | 0.89 | 0.92 (0.08-10.6) | 0.94 |
|  | Endometrioid | 1.69 (0.93-3.08) | 0.08 | 1.09 (0.11-11.2) | 0.94 | 1.00 (0.98-1.03) | 0.70 | 1.95 (0.84-4.55) | 0.12 | 0.67 (0.08-5.70) | 0.71 |
|  | Clear cell | 0.86 (0.38-1.92) | 0.71 | 2.68 (0.11-62.9) | 0.54 | 0.99 (0.96-1.02) | 0.46 | 0.71 (0.22-2.28) | 0.56 | 0.37 (0.01-12.5) | 0.58 |
|  | LMP | 1.25 (0.84-1.85) | 0.26 | 0.85 (0.19-3.81) | 0.83 | 1.01 (0.98-1.03) | 0.60 | 1.39 (0.78-2.47) | 0.27 | 0.98 (0.23-4.15) | 0.97 |

Causal estimates are scaled to represent the effect of a one-unit increase in natural log 25-hydroxyvitamin D (ng/mL) and a one-unit increase in comprehensive smoking exposure. IVW = Inverse-variance weighted, HGSC = High grade serous carcinoma, LGSC = Low grade serous carcinoma, LMP = Low malignant potential.

**Supplementary Table 6. IVW and sensitivity analysis estimates for causal estimates of molecular risk factors on ovarian cancer risk**

Causal estimates are scaled to represent the effect of a 1-unit increase in natural log-transformed CRP (mg/L) and a 1-unit increase in natural log-transformed sex hormone-binding globulin (nmol/L). IVW = Inverse-variance weighted, HGSC = High grade serous carcinoma, LGSC = Low grade serous carcinoma, LMP = Low malignant potential.

| **Risk factor** | **Ovarian cancer outcome** | **IVW**  **OR (95% CI)** | ***P*-value** | **MR-Egger regression**  **OR (95% CI)** | ***P*-value** | **MR-Egger intercept**  **OR (95% CI)** | ***P*-value** | **Weighted median**  **OR (95%CI)** | ***P*-value** | **Weighted mode**  **OR (95% CI)** | ***P*-value** |
| --- | --- | --- | --- | --- | --- | --- | --- | --- | --- | --- | --- |
| **C-reactive protein** | | | | | | | | | | | |
|  | Overall | 0.97 (0.93-1.02) | 0.19 | 0.99 (0.93-1.06) | 0.86 | 0.99 (0.98-1.01) | 0.37 | 0.98 (0.93-1.03) | 0.42 | 0.98 (0.93-1.03) | 0.41 |
|  | HGSC | 0.99 (0.93-1.05) | 0.66 | 1.03 (0.94-1.11) | 0.58 | 0.99 (0.97-1.01) | 0.26 | 0.99 (0.93-1.05) | 0.65 | 0.99 (0.93-1.06) | 0.86 |
|  | LGSC | 0.89 (0.69-1.14) | 0.36 | 0.79 (0.54-1.15) | 0.27 | 1.04 (0.95-1.12) | 0.43 | 0.88 (0.73-1.05) | 0.15 | 0.88 (0.73-1.06) | 0.21 |
|  | Mucinous | 0.90 (0.78-1.04) | 0.14 | 0.92 (0.74-1.14) | 0.46 | 0.99 (0.95-1.04) | 0.82 | 0.91 (0.78-1.07) | 0.25 | 0.91 (0.77-1.07) | 0.27 |
|  | Endometrioid | 0.90 (0.82-1.00) | 0.049 | 0.96 (0.83-1.11) | 0.59 | 0.99 (0.95-1.04) | 0.82 | 0.93 (0.83-1.03) | 0.18 | 0.93 (0.83-1.03) | 0.20 |
|  | Clear cell | 1.00 (0.87-1.16) | 0.96 | 0.96 (0.78-1.18) | 0.69 | 1.01 (0.97-1.06) | 0.55 | 0.98 (0.84-1.13) | 0.76 | 0.97 (0.83-1.13) | 0.71 |
|  | LMP | 0.99 (0.89-1.11) | 0.86 | 1.01 (0.85-1.20) | 0.93 | 0.99 (0.96-1.03) | 0.79 | 1.01 (0.91-1.13) | 0.81 | 0.99 (0.89-1.10) | 0.86 |
| **Sex hormone-binding globulin** | | | | | | | | | | | |
|  | Overall | 0.98 (0.78-1.23) | 0.85 | 1.47 (0.80-2.71) | 0.27 | 0.98 (0.95-1.01) | 0.21 | 1.09 (0.83-1.45) | 0.53 | 1.14 (0.83-1.56) | 0.45 |
|  | HGSC | 1.08 (0.83-1.42) | 0.55 | 1.15 (0.56-2.37) | 0.72 | 1.00 (0.96-1.03) | 0.87 | 1.09 (0.78-1.53) | 0.61 | 1.10 (0.75-1.63) | 0.64 |
|  | LGSC | 0.84 (0.24-2.98) | 0.79 | 4.99 (0.18-136.5) | 0.38 | 0.91 (0.78-1.07) | 0.31 | 1.17 (0.39-3.52) | 0.78 | 1.49 (0.49-4.59) | 0.51 |
|  | Mucinous | 0.64 (0.33-1.26) | 0.19 | 1.06 (0.17-6.49) | 0.95 | 0.97 (0.89-1.06) | 0.58 | 0.75 (0.32-1.76) | 0.50 | 0.76 (0.29-2.01) | 0.60 |
|  | Endometrioid | 0.77 (0.36-1.62) | 0.49 | 1.00 (0.11-8.99) | 0.99 | 0.99 (0.89-1.10) | 0.81 | 0.99 (0.51-1.92) | 0.97 | 1.10 (0.54-2.23) | 0.81 |
|  | Clear cell | 1.03 (0.52-2.04) | 0.93 | 4.85 (0.78-30.1) | 0.15 | 0.92 (0.85-1.01) | 0.13 | 1.55 (0.64-3.72) | 0.33 | 1.80 (0.67-4.84) | 0.29 |
|  | LMP | 1.00 (0.58-1.74) | 0.99 | 1.01 (0.30-3.42) | 0.98 | 1.00 (0.94-1.06) | 0.98 | 0.96 (0.55-1.67) | 0.88 | 0.97 (0.55-1.72) | 0.92 |
